# Xylazine Reshapes the Sleep and Respiratory Consequences of Chronic Fentanyl Exposure in a Sex-Dependent Manner

**DOI:** 10.64898/2026.09.02.748951

**Authors:** Mackenzie C. Gamble, Samara J. Vilca, Benjamin R. Williams, Grace Guidon, Vivian Dang, Ryan W. Logan

## Abstract

Recent shifts in the illicit fentanyl supply have led to widespread exposure to xylazine, yet the biological consequences of chronic fentanyl-xylazine co-exposure remain poorly understood. Here, we compared the behavioral and molecular effects of chronic exposures to fentanyl and xylazine both alone and in combination in male and female mice. Combined fentanyl-xylazine produced distinct, sex-dependent alterations in sleep architecture, including changes in NREMS recovery, REMS dynamics, and reduced sleep bout duration compared with fentanyl alone. The α_2_-adrenergic receptor antagonist yohimbine partially reversed these sleep alterations in a sex-dependent manner. Fentanyl-xylazine also exacerbated respiratory depression relative to fentanyl alone, particularly in males. In contrast, despite its prevalence in humans no skin lesions were observed. Molecular analyses revealed that fentanyl-xylazine co-administration leads to increased peripheral Fibroblast growth factor 21, a hepatokine linked to reduced opioid preference and alcohol consumption. Together, these findings demonstrate that xylazine substantially alters the physiological and neurobiological consequences of chronic fentanyl exposure in a sex-dependent manner, providing a preclinical framework for understanding the growing public health impact of fentanyl-xylazine co-use.

## Introduction

Fentanyl and its analogs remain the leading cause of overdose deaths (Cano et al., 2023); however, recent changes in the drug supply have resulted in large increases in alpha 2 adrenergic agonists (α_2_ agonists) in fentanyl samples. Specifically, evidence of fentanyl adulteration with the α_2_ agonist xylazine began appearing around 2020 (Marshall and Nelson, 2025). Xylazine is a veterinary tranquilizer not approved for human use globally. While xylazine is a recent and mostly unintentional component of addiction, α_2_ agonists have a documented history of intentional co-use with opioids. For instance, individuals have purposefully used clonidine in combination with opioids for its psychoactive effects (Beuger et al., 1998; Dennison, 2001; Mitchell and Lee, 2021). However, the emergence of xylazine in the illicit drug supply has been associated with distinct clinical effects that have not been observed with clonidine.

Studies have described a range of distinct and severe adverse effects associated with concomitant fentanyl-xylazine use. Individuals exposed to fentanyl-xylazine can develop severe necrotic skin wounds, that in extreme cases can result in amputation (Arango et al., 2024; Laurano et al., 2025; Oscherwitz et al., 2024). Additionally, the combination of these drugs appears to complicate opioid overdose responses, increasing the risk of potentially lethal respiratory depression (Demery et al., 2025a). Reported opioid withdrawal symptoms are also altered, with emerging evidence indicating atypical patterns such as fluctuations in heart rate and heightened anxiety in individuals exposed to xylazine (Alexander et al., 2025; Thakrar et al., 2025). Xylazine exposure has also been associated with prolonged sedation. Qualitative data indicate that it may provoke parasomnias, suggesting distinct sleep-related disturbances unique to fentanyl-xylazine use, which are not observed with fentanyl alone (Reed et al., 2025, 2022). At high doses, most α_2_ agonists increase light non-rapid eye movement sleep (NREMS, N1-2) and decrease REMS (Chamadia et al., 2020; Garrity et al., 2015; Seidel et al., 1995; Spiegel and DeVos, 1980). However, clonidine has opposing effects depending on dose. Low-dose clonidine tends to increase REMS while decreasing NREMS and vice versa at higher doses in humans (Miyazaki et al., 2004). Yet, the sleep and withdrawal-related effects of subanesthetic doses of xylazine, whether they are administered on their own or with fentanyl, are unknown.

Overall, the sudden proliferation of xylazine has led to a significant information gap regarding the α_2_-adrenergic system during opioid addiction. Many of its effects have yet to be carefully characterized, and some studies offer conflicting pictures of the effects of xylazine adulteration (Hill et al., 2025; Perrone et al., 2024; Sibley et al., 2025; Tan et al., 2024). Despite the broad range of reported effects, few studies have systematically investigated these observations or their underlying mechanisms. Thus, the goal of this study is to provide preclinical characterization of fentanyl-xylazine’s effects across four domains including: respiration, sleep-wake, withdrawal, and molecular changes.

## Methods

### Animals and Housing

168 adult (9–15-week-old) male and female C57BL/6J mice (Jackson Laboratory Strain #000664) were used in total. The cohorts were balanced by sex with the following sample sizes according to the respective endpoints: sleep-wake analysis (n=64 with n=8 for the yohimbine arm), respiratory analysis (n=64), and molecular analysis (n=40). Mice were group housed in a 12:12 light-dark cycle (lights on at 0700h, ZT0, and off at 1900h, ZT12) with standard rodent chow and water provided *ad libitum*. For sleep-wake and respiratory cohorts, mice were transferred from group-housed home cages to single-housing in PiezoSleep Mouse Behavioral Tracking Systems (Signal Solutions, Inc., Lexington, KY, USA), maintained at the same 12:12 light-dark cycle, and given several days to acclimate. For molecular cohorts, mice were group-housed throughout the experiment. All experimental procedures were approved by the Institutional Animal Care and Use Committee at University of Massachusetts Chan Medical School (IPROTO202300000010).

### Drugs

All drugs were dissolved in 0.9% sterile filtered saline. Fentanyl citrate solution (50 μg/mL, Patterson Veterinary Supply, Inc., Mt. Joy, PA, USA) and xylazine hydrochloride (Millipore Sigma, St. Louis, MO, USA) were administered by intraperitoneal injection (i.p.). For sleep-wake, withdrawal, and molecular cohorts, fentanyl and xylazine hydrochloride was dosed at 320 μg/kg and 5 mg/kg, respectively, twice daily at ZT1 and ZT9. For the respiratory cohort, the xylazine dose was kept the same throughout, while fentanyl dosing was as follows: 0 μg/kg, 320 μg/kg, 640 μg/kg, and 1.28 mg/kg. Xylazine dose was based on previous xylazine polysubstance studies and designed to be subanesthetic. Yohimbine hydrochloride (Millipore Sigma) was injected (i.p.) at ZT1 only at 1.5 mg/kg. Yohimbine dose was based on previous dosing shown to reverse xylazine anesthesia.

### Experimental Paradigms

Figure 1 outlines the experimental details of all cohorts. For the sleep-wake cohort, mice were habituated to recording chambers (Signal Solutions, Inc., Lexington, KY, USA) for 5-7 days. After habituation, a single baseline day was recorded (mice undisturbed). Following the baseline recording, drugs (saline, xylazine, fentanyl, or fentanyl-xylazine) were administered twice daily, ∼8 hours apart (ZT1, 0800h and ZT9, 1600h), for 7 days. Yohimbine hydrochloride was only administered once daily at ZT1. After 7 days, mice were allowed to undergo acute withdrawal (24 hours with no drug). Half of all mice were pseudo-randomly assigned for withdrawal recording including baseline and acute withdrawal. For the respiratory cohort, mice were habituated to the recording chambers for at least 48 hours. After 48 hours, 24 hours of baseline recording were performed followed by administration of a single dose of fentanyl with or without a fixed dose of xylazine (the zero-fentanyl conditions being saline and xylazine alone, respectively) in naive mice. Lastly, for the molecular analysis cohort, mice were group-housed in home cages and treated at ZT1 and ZT9 for seven days, as in the sleep-wake cohort. Mice were then euthanized, and tissue was collected approximately two hours after final ZT1 injection on the 7^th^ day.

**Figure 1.**
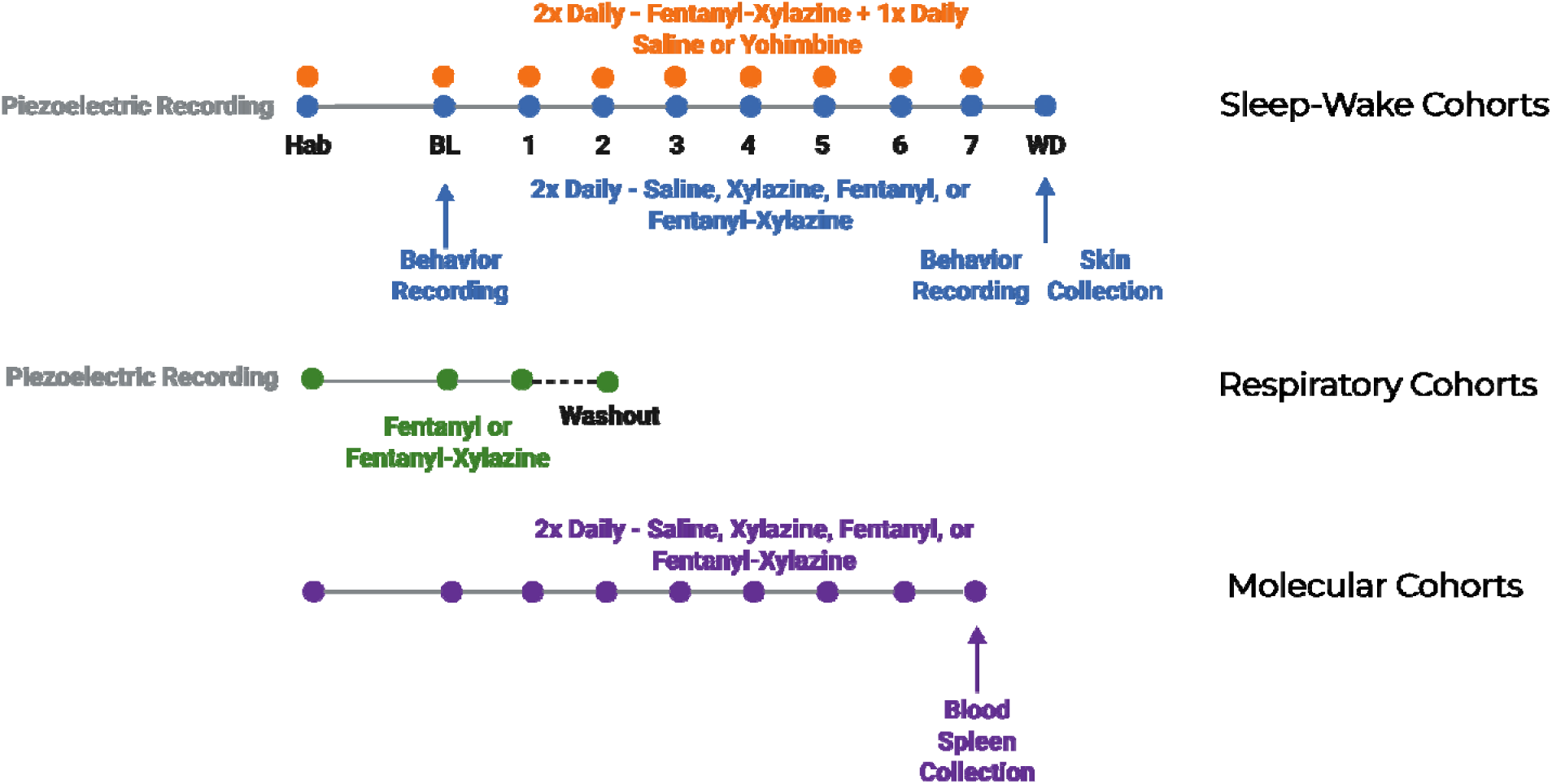
Experimental paradigms of different fentanyl-xylazine cohorts.

### Piezoelectric Recordings

During habituation and recording, mice were housed in individual 6-inch x 6-inch polycarbonate cages (PiezoSleep Mouse Behavioral Tracking Systems,Lexington, KY, USA) with piezoelectric sensor floors attached to a computer. Activity signals (both macro and micro) were recorded and categorized as Wake, NREMS, or REMS using SleepStats software version 4 (Signal Solutions, Inc., Lexington, KY, USA), binned in 1-hour intervals, then averaged per 4 hours. Respiratory features were extracted using custom software in collaboration with Signal Solutions. Briefly, the software collects 24-hour baseline of average respiration and then detects any drop in respiration greater than 10% from baseline to determine a respiratory depression event and its related features. Injection times are inputted to time lock analyses.

### Withdrawal Recording and Scoring

Baseline and withdrawal behaviors were recorded in a transparent plexiglass tube using a web camera (720p) ∼18 hours after the final injection of drug. A thin layer of bedding was provided and mice recorded individually for 15 min. Behaviors included rearing (supported or unsupported), grooming (face or body), wet-dog shakes, digging (targeted movement of bedding), and scratching (rear paw on body or head). Other classic opioid withdrawal-related behaviors like ptosis, teeth chattering, piloerection, jumping, freezing, jaw/paw tremor and diarrhea were assessed but not observed. Each mouse was scored by two separate individuals and an average score per mouse compiled for analysis.

### Tissue Collection, RNA Extraction, and qPCR

Blood, brain, spleen, and skin were extracted from each animal. Blood was collected from the trunk via rapid decapitation, let clot, and then centrifuged at 2,000 x g for 15 min at 4°C to extract serum. Brain and spleen were flash frozen in isopentane chilled on dry ice, while skin from the mouse paw pad was placed in TRIzol Reagent (Life Technologies, Logan, UT, USA) and frozen directly on dry ice. All tissues were stored at −80°C before being processed. RNA extraction from mouse skin and spleen was performed using Dounce homogenization (KIMBLE Dounce tissue grinder, Millipore Sigma) and RNeasy Mini Kits (Qiagen Inc., Germantown, MD, USA). The homogenate was processed through a QIAshredder column (Qiagen Inc., Germantown, MD, USA) to ensure complete homogenization. RNA was then purified using the RNeasy Mini Kit, including an on-column DNase I digestion step. RNA concentration and purity were assessed using a NanoDrop spectrophotometer (Thermo Fisher Scientific LLC, Asheville, NC, USA). For qPCR, Bio-Rad primers (Bio-Rad Laboratories, Hercules, CA, USA) were used for target genes and housekeeping genes (GAPDH). Reactions were run in duplicate. Relative gene expression was calculated using the 2^(−ΔΔCt) method and statistics were performed on ΔΔCt values.

### Proximity Extension Assay

Mouse serum (40 samples; balanced by treatment group, cage, and sex) was processed via targeted proteomics using the highly sensitive Olink proximity extension assay (Olink Proteomics Inc., Waltham, MA, USA). Briefly, Olink detects the presence of proteins using oligonucleotide-linked antibodies coupled with PCR to turn protein detection into a quantifiable signal. The mouse Olink Target 48 immune panel was used which detects 45 cytokines, chemokines, and other immune regulatory proteins.

### Statistics

Data were analyzed using ANOVA (one-way ordinary, two-way ordinary and two-way repeated measures) via GraphPad Prism or linear mixed-effects models in R. One-way ordinary or Welch ANOVA was used for circadian and rhythm metrics, two-way ordinary for respiration metrics, and two-way repeated measures for sleep metrics. Holm post-hoc corrections were used for all multiple comparisons. Additionally, withdrawal behaviors were analyzed using linear mixed-effects models (lmer package), with time (to account for baseline recordings) and treatment as fixed effects, and Animal ID included as a random intercept to account for repeated measures. Low-frequency, zero-inflated behavioral measures (average shaking and scratching counts) were log1p-transformed prior to analysis. Olink data were also analyzed using linear mixed-effects models were fitted with drug exposure as a fixed effect and cage as a random intercept to account for shared environmental/housing effects among animals within the same cage. Proteins meeting the significance threshold from the mixed-effects model analysis (FDR corrected) were further evaluated using post hoc pairwise condition comparisons with Tukey correction using the OlinkAnalyze R package. Results are presented as mean ± standard error of the mean (SEM), with significance set at α=0.05

## Results

We characterized the respective effects of saline, subanesthetic xylazine, fentanyl, or an adulteration of subanesthetic xylazine with fentanyl (2x daily at ZT 1 and 9) on circadian activity and vigilance states. These data were analyzed in both sexes and averaged across seven days (Figure 1). For simplicity, vigilance state results are described across the light-dark period with analyses per 3 ZTs noted when relevant.

### Fentanyl-Only Decreases Interdaily Stability in Females

In both female and male mice, there were no discernable differences in percentage of dark phase activity (female: F (3, 15.58) = 0.2430, P = 0.8650; male: F (3,20) = 2.537, P = 0.0857), relative amplitude of activity (female: F (3,20) = 2.174, P = 0.1228; male: F (3,20) = 1.924, P = 0.1582), and intradaily variability (female: F (3,20) = 0.2039, P = 0.8925; male: F (3,20) = 1.547, P = 0.2333) across treatment groups over seven days of administration (Figure 2 A-C & E-G). Interdaily stability was, however, significantly different in females (F (3,20) = 3.665, P = 0.0297), but not males (F (3,10.04) = 3.289, P = 0.0663). Post-hoc analysis indicated that interdaily stability in fentanyl-treated female mice was significantly decreased compared to saline (P = 0.0375), but not between other treatment groups (Figure 2D).

**Figure 2.**
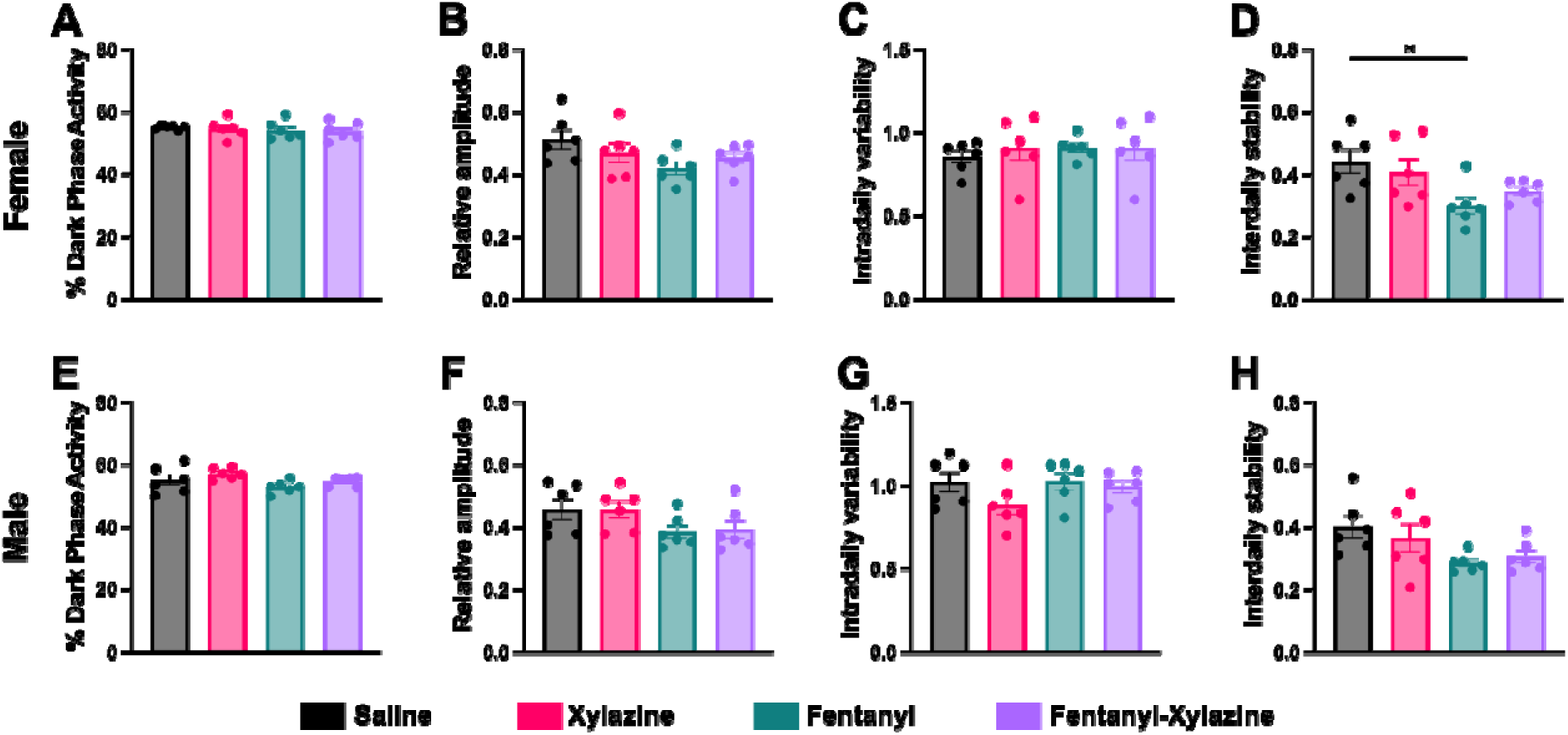
Alterations in circadian activity and rhythm structure from fentanyl and fentanyl-xylazine administration in female and male mice. **A** Percent dark phase activity in females. **B** Relative amplitude of activity in females. **C** Intradaily variability in females. **D** Intradaily stability in females. **E** Percent dark phase activity in males. **F** Relative amplitude of activity in males. **G** Intradaily variability in males. **H** Intradaily stability in males. Data presented as mean ± SEM; *p < 0.05.

### Co-administered Fentanyl-Xylazine Mirrors Fentanyl Sleep Disruption in Females but Modifies Dark Period NREMS Recovery in Males

In both female and male mice, two-way repeated measures ANOVA revealed a significant main effect of light-dark phase (female: F (1, 20) = 46.52, P < 0.0001; male: F (1, 20) = 12.47, P = 0.0021) and a significant treatment × light-dark phase interaction (female: F (3, 20) = 11.17, P = 0.0002; male: F (3, 20) = 49.68, P < 0.0001) for wakefulness (Figure 3A&B). A significant main effect of treatment was observed in males (F (3, 20) = 3.979, P = 0.0225), but not in females (F (3, 20) = 1.378, P = 0.2784).

**Figure 3.**
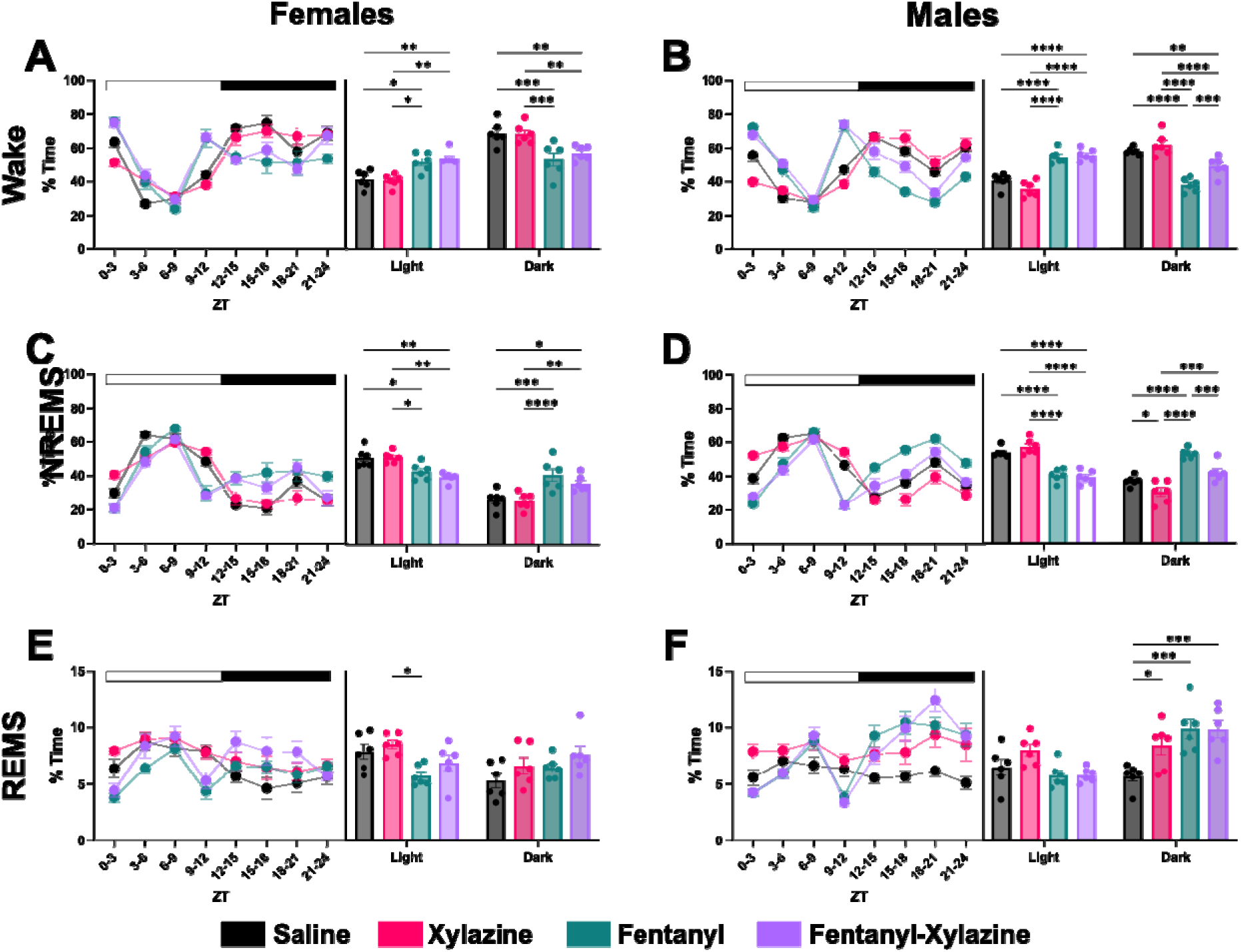
Alterations in the percent time across vigilance states from fentanyl and fentanyl-xylazine administration in female and male mice. **A** Time spent awake across seven days of recording expressed as in tri-ZT and light/dark bins in females. **B** Time spent awake across seven days of recording expressed as in tri-ZT and light/dark bins in males. **C** Time spent in NREMS across seven days of recording expressed as in tri-ZT and light/dark bins in females. **D** Time spent in NREMS across seven days of recording expressed as in tri-ZT and light/dark bins in males. **E** Time spent in REMS across seven days of recording expressed as in tri-ZT and light/dark bins in females. **F** Time spent in REMS across seven days of recording expressed as in tri-ZT and light/dark bins in males. Data presented as mean ± SEM; *p < 0.05, **p < 0.01, ***p < 0.001, ****p < 0.0001.

Holm-corrected post hoc testing identified significant differences between treatments within each light-dark phase for each sex. During the light period, both fentanyl and fentanyl-xylazine-treated mice exhibited similar levels of wakefulness in females (P = 0.7623) and males (P = 0.7071). In both sexes, wake time in fentanyl and fentanyl-xylazine groups was significantly elevated compared to saline (Females: P = 0.0184 & P = 0.0050; Males: P < 0.0001 & P < 0.0001) and xylazine (Females: P = 0.0128 & P = 0.0029; Males: P < 0.0001 & P < 0.0001) controls, which themselves did not differ from each other (P = 0.8098; P = 0.137) in either sex.

In contrast, during the dark period, both fentanyl and fentanyl-xylazine led to significantly less time spent awake relative to saline and xylazine groups in both female (P = 0.0004 & P = 0.0071; P = 0.0005 & P = 0.0073) and male (P < 0.0001 & P = 0.003; P < 0.0001 & P < 0.0001) mice. Among males, fentanyl administration reduced wakefulness compared to fentanyl-xylazine together (P = 0.0005). No such difference was observed between fentanyl and fentanyl-xylazine exposure in females (P = 0.4742). Additionally, saline and xylazine-treated mice did not differ in dark-period wakefulness in either sex (P = 0.9096, P = 0.1707).

In both female and male mice, two-way repeated measures ANOVA revealed there was a significant main effect of treatment (female: F (3, 20) = 3.33, P = 0.0403; male: F (3, 20) = 4.9830, P = 0.0096), light-dark phase (female: F (1, 20) = 51.72, P < 0.0001; male: F (1, 20) = 27.86, P < 0.0001), and interaction (female: F (3, 20) = 10.77, P = 0.0002; male: F (3, 20) = 45.29, P < 0.0001) for NREMS (Figure 3 C&D). Holm-corrected post-hoc testing indicated significant differences within in each light-dark phase between treatments for each sex, respectively. During the light phase, the increase in wakefulness following fentanyl and fentanyl-xylazine exposure was mirrored by a significant decrease in NREMS for both sexes, as compared to saline (females: P = 0.05 & P = 0.0037, males: P < 0.0001 & P < 0.0001) and xylazine (females: P = 0.05 & P = 0.0037, males: P < 0.0001 & P < 0.0001) groups. No significant differences were found between saline and xylazine (P = 0.95, P = 0.319), nor between fentanyl and fentanyl-xylazine (P = 0.475, P = 0.6798) within either sex.

During the dark period, patterns diverged by sex. In female mice, fentanyl and fentanyl-xylazine administration resulted in significantly greater NREMS compared to both saline (P = 0.0002, P = 0.0138) and xylazine (P < 0.0001, P = 0.0083) controls. No differences in dark-period NREMS were noted between saline and xylazine (P = 0.7727), nor between fentanyl and fentanyl-xylazine (P = 0.2157) in females. In male mice, fentanyl increased NREMS compared to saline (P < 0.0001), while fentanyl-xylazine did not differ from saline (P = 0.0533). Further, fentanyl-treated males spent more time in NREMS than those receiving fentanyl-xylazine (P = 0.0002), and xylazine-treated males exhibited reduced NREMS relative to saline (P = 0.026). Both fentanyl (P < 0.0001) and fentanyl-xylazine (P = 0.0002) treated males had significantly higher NREMS than those treated with xylazine alone.

In both female and male mice, two-way repeated measures ANOVA revealed the absence of a main effect of treatment (female: F (3, 20) = 1.593, P = 0.2225; male: F (3, 20) = 2.464 P = 0.0921). Instead, there was a significant main effect of light-dark phase (female: F (1, 20) = 6.228, P = 0.0214; male: F (1, 20) = 72.87, P < 0.0001) and interaction (female: F (3, 20) = 8.083, P = 0.001; male: F (3, 20) = 30.37, P < 0.0001) for REMS (Figure 3 E&F). Holm-corrected post-hoc testing indicated significant differences within the dark phase only for males and light phase only for females. During the light period, only female fentanyl-treated mice spent significantly less time in REMS than female xylazine-treated mice (P = 0.0145). REMS was slightly reduced in both fentanyl and fentanyl-xylazine groups compared to saline and xylazine groups; however, these differences were non-significant No other significant differences were observed between any female groups, and treatment groups did not differ in males during the light phase.

In the dark period, there were no significant differences in REMS among all female groups including saline, xylazine, fentanyl, and fentanyl-xylazine. Conversely, in males, xylazine (P = 0.0191), fentanyl (P = 0.0003), and fentanyl-xylazine (P = 0.0003) exposure each resulted in significantly greater REMS compared to saline. In females, fentanyl and its combination with subanesthetic xylazine produced comparable alterations in all three vigilance states across both the light and dark periods, with the combination reflecting the disruptive effects of fentanyl alone. In males, while fentanyl and fentanyl-xylazine produced similar patterns during the light period, their effects diverged in the dark period: fentanyl-xylazine-treated males had wake and NREMS amounts more closely matching saline controls, while fentanyl alone produced the greatest increase in wake and thus in-kind decrease in NREMS. The elevation of REMS during the dark phase was a unique effect observed only in males following xylazine, fentanyl, or their combination.

### Fentanyl-Xylazine Attenuates Wake Bout Duration and Reduces NREMS Bout Length Across Sexes Compared to Fentanyl During the Light Period

In female mice, two-way repeated measures ANOVA revealed there was a significant main effect of treatment (F (3, 20) = 4.587, P = 0.0134), light-dark phase (F (1, 20) = 13.58, P = 0.0015), and interaction (F (3, 20) = 5.909, P = 0.0047). In males there was a significant main effect of treatment (F (3, 20) = 4.309, P = 0.0169) and interaction effect (F (3, 20) = 14.49, P < 0.0001), but not of light-dark phase (F (1, 20) = 3.137, P = 0.0918) for average wake bout duration (Figure 4 A&B). Holm-corrected post-hoc testing indicated during the light period, fentanyl led to significantly elevated average wake bout durations in comparison to xylazine-only in females (P = 0.0288) and compared to saline, xylazine, and fentanyl-xylazine in males (saline: P < 0.0001, xylazine: P < 0.0001, fentanyl-xylazine: P = 0.0005). Looking by ZT, it is clear that the first injection of fentanyl is comparable between females and males, but the second injection does not lead to as robust of a longer wake bout response in females compared to males. Fentanyl-xylazine largely normalized this effect with no difference between fentanyl-xylazine and saline in females (P = 0.6741) and males (P = 0.3924). There were no differences between each of the other groups.

**Figure 4.**
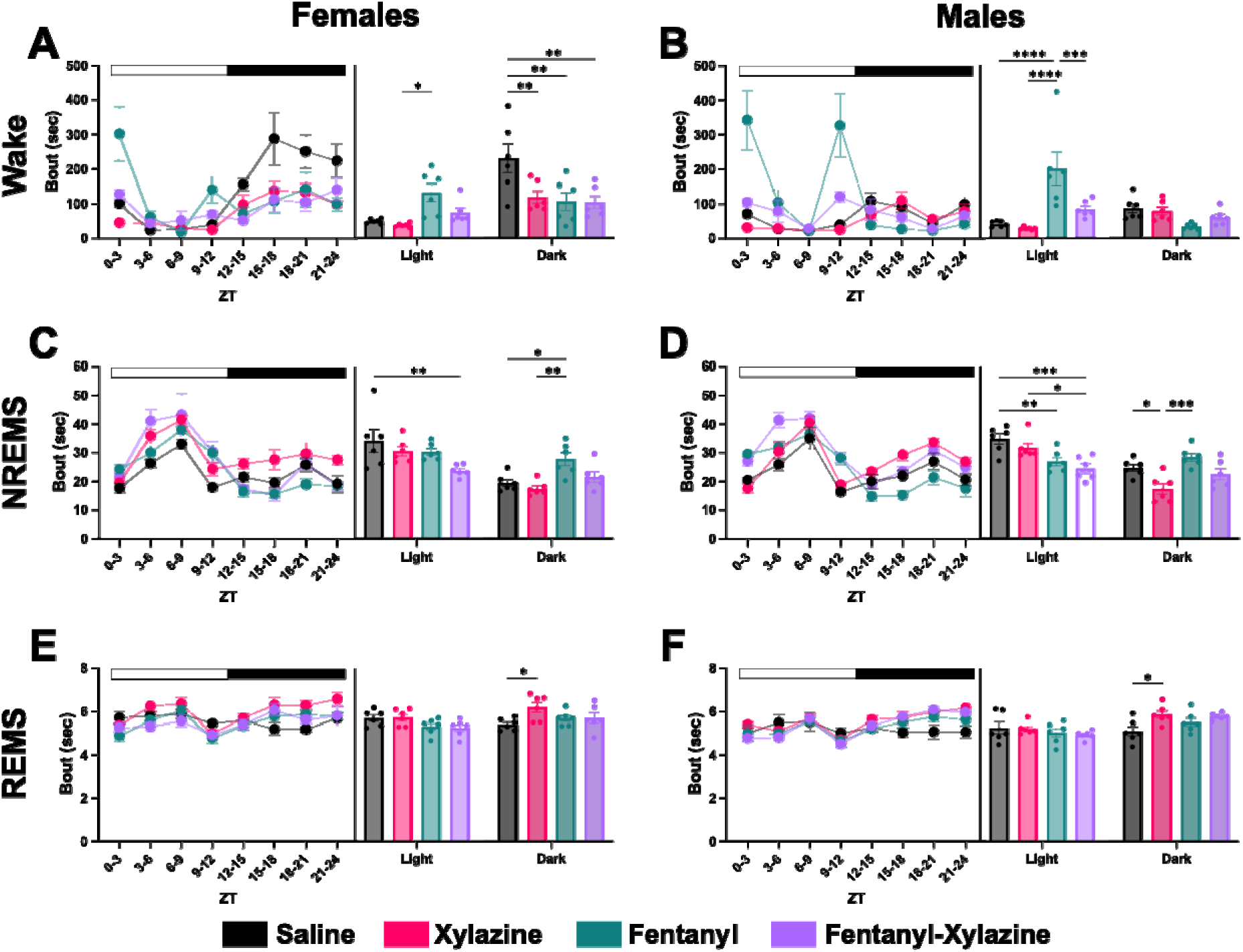
Alterations in the bout durations across vigilance states from fentanyl and fentanyl-xylazine administration in female and male mice. **A** Average wake bout duration across seven days of recording expressed as in tri-ZT and light/dark bins in females. **B** Average wake bout duration across seven days of recording expressed as in tri-ZT and light/dark bins in males. **C** Average NREMS bout duration across seven days of recording expressed as in tri-ZT and light/dark bins in females. **D** Average NREMS bout duration across seven days of recording expressed as in tri-ZT and light/dark bins in males. **E** Average REMS bout duration across seven days of recording expressed as in tri-ZT and light/dark bins in females. **F** Average REMS bout duration across seven days of recording expressed as in tri-ZT and light/dark bins in males. Data presented as mean ± SEM; *p < 0.05, **p < 0.01, ***p < 0.001, ****p < 0.0001.

During the dark period, female saline-treated mice exhibited higher average wake bout duration than xylazine (P = 0.0042), fentanyl (P = 0.0017), and fentanyl-xylazine (P = 0.0016) treated females. All other groups were comparable. In males, no significant differences between any groups were noted in the dark period.

In both female and male mice, two-way repeated measures ANOVA revealed there was a significant main effect of treatment (female: F (3, 20) = 4.029, P = 0.0215; male: F (3, 20) = 4.132, P = 0.0197), light-dark phase (female: F (1, 20) = 31.85, P<0.0001; male: F (1, 20) = 46.66, P < 0.0001), and interaction (female: F (3, 20) = 5.374, P = 0.0071; male: F (3, 20) = 15.69, P < 0.0001) for average NREMS bout duration (Figure 4 C&D). Interestingly, average NREMS bout duration was significantly lower in males and females administered fentanyl-xylazine compared to saline (females: P = 0.0051, males: P = 0.0006) in the light period. While the fentanyl-xylazine group had the lowest NREMS bout duration than all other groups in both sexes, there were no significant differences between fentanyl and fentanyl-xylazine (females: P = 0.1069, males: P = 0.3504). Additionally, in males only, fentanyl significantly reduced NREMS bout duration compared to saline (P = 0.0098) and likewise fentanyl-xylazine administration reduced NREMS bout duration compared to xylazine (P = 0.0187).

In the dark period, fentanyl-treated mice had the longest average NREMS bout duration in both sexes. In females, this was significantly higher compared to saline (P = 0.028) and xylazine (P = 0.005) groups, but not fentanyl-xylazine (P = 0.1329). In males, this was only significantly higher than xylazine (P = 0.0002), which was significantly lower than saline treated males (P = 0.0187). No differences were noted between fentanyl and fentanyl-xylazine.

In both female and male mice, two-way repeated measures ANOVA revealed there was no main effect of treatment (female: F (3, 20) = 2.138, P = 0.1274; male: F (3, 20) = 0.7899, P = 0.5137), but a significant main effect of light-dark phase (female: F (1, 20) = 10.10, P = 0.0047; male: F (1, 20) = 74.84, P < 0.0001) and interaction effect (female: F (3, 20) = 5.307, P = 0.0074; male: F (3, 20) = 16.77, P < 0.0001) for average REMS bout duration (Figure 4 E&F). Notably, Holm-corrected post-hoc testing indicated no differences between any groups in both sexes in the light period. However, in the dark period, xylazine-treated mice had the highest average REMS bout duration in both sexes which was only significant compared to saline-treated mice (female: P = 0.0163, male: P = 0.0295). No other differences were noted between groups in either sex.

In summary, fentanyl-xylazine-treated male and female mice had reduced average wake bout durations at the time of injections (ZT 1 & 9, light period) and more closely resembled saline-treated mice compared to the fentanyl-treated group. Additionally, fentanyl-xylazine-treated mice had the lowest average NREMS bout duration during the same period across sexes. Fentanyl-treated mice experienced NREMS bout duration rebound in the dark period following increased wake bout duration in the light period, again in both sexes. Xylazine uniquely increased REMS bout duration in the dark period in both sexes.

### Yohimbine Differentially Negates Fentanyl-Xylazine-Induced Wake and NREMS Changes in Males and REMS Suppression in Females

Two-way repeated measures ANOVA indicated there were no significant differences due to treatment (F (1, 6) = 0.2887, P = 0.6104), light-dark phase (F (1, 6) = 5.381, P = 0.0595), or interaction (F (1, 6) = 0.0085, P = 0.9294) in females for wake (Figure 5A). Thus, the combination of yohimbine and fentanyl-xylazine produced no significant alterations in time spent in wake compared to fentanyl-xylazine alone in females. A closer look by ZT reaches the same conclusion. In males, there was only a main effect of treatment (F (1, 6) = 6.867, P = 0.0396) and not light-dark phase (F (1, 6) = 1.10, P = 0.3347) or an interaction effect (F (1, 6) = 0.3508, P = 0.5753) for wake (Figure 5B). The combination of yohimbine with fentanyl-xylazine lowered time spent in wake compared to vehicle irrespective of light-dark phase. Looking closer by ZT, this effect was largely more concentrated to the later portion of the dark period.

**Figure 5.**
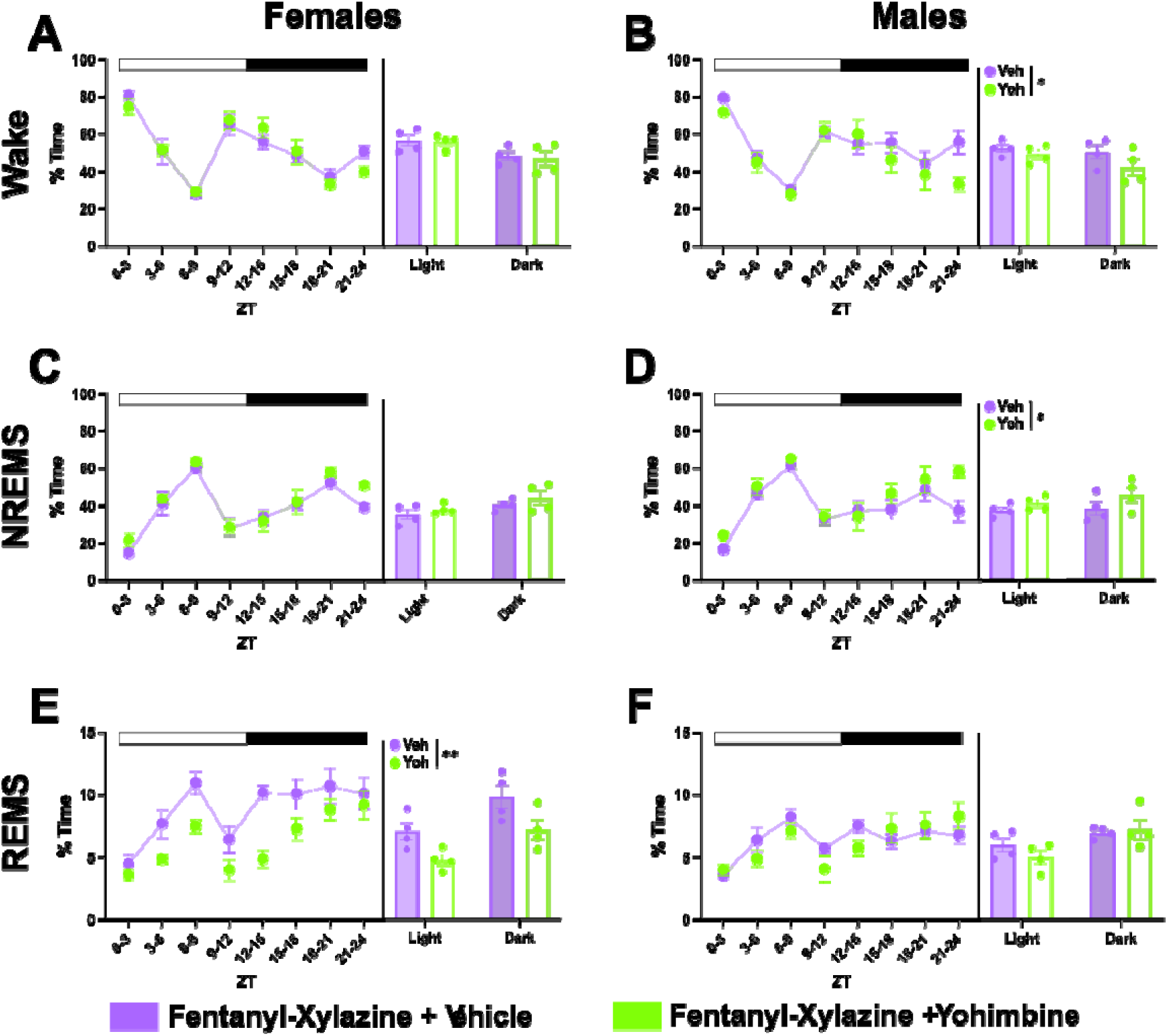
Alterations in the percent time across vigilance states from yohimbine treatment in female and male mice. **A** Time spent awake across seven days of recording expressed as in tri-ZT and light/dark bins in females. **B** Time spent awake across seven days of recording expressed as in tri-ZT and light/dark bins in males. **C** Time spent in NREMS across seven days of recording expressed as in tri-ZT and light/dark bins in females. **D** Time spent in NREMS across seven days of recording expressed as in tri-ZT and light/dark bins in males. **E** Time spent in REMS across seven days of recording expressed as in tri-ZT and light/dark bins in females. **F** Time spent in REMS across seven days of recording expressed as in tri-ZT and light/dark bins in males. Data presented as mean ± SEM; *p < 0.05, **p < 0.01.

In line with no differences detected between yohimbine-treated and control female mice for wake, two-way repeated measures ANOVA indicated no main effects (treatment: F (1, 6) = 3.254, P = 0.1213, light-dark phase: F (1, 6) = 3.699, P = 0.1028) or interaction effect (F (1, 6) = 0.0269, P = 0.8751) (Figure 5C). However, males exhibited a significant main effect of treatment again (F (1, 6) = 6.495, P = 0.0436) and no main effect of light-dark phase (F (1, 6) = 0.6199, P = 0.4611) or interaction effect (F (1, 6) = 0.2666, P = 0.6241) (Figure 5D). There was a concordant increase in time spent in NREMS in fentanyl-xylazine-yohimbine-treated male mice compared to fentanyl-xylazine-treated mice irrespective of light dark phase.

Two-way repeated measures ANOVA indicated a significant main effect of treatment (F (1, 6) = 14.04, P = 0.0095) and light-dark phase (F (1, 6) = 10.20, P = 0.0187), but no interaction effect (F (1, 6) = 0.0346, P = 0.8585) in females for REMS (Figure 5E). Notably, fentanyl-xylazine-yohimbine administration led to decreased time spent in REMS compared to fentanyl-xylazine alone in females regardless of light-dark phase. However, no differences in REMS between groups were noted in males with two-way repeated measures ANOVA, indicating no main effect of treatment (F (1, 6) = 0.5281 P = 0.4948), light-dark phase (F (1, 6) = 5.848, P = 0.052), or interaction (F (1, 6) = 0.9051, P = 0.3781) (Figure 5F).

In summary, the combination of once daily yohimbine with fentanyl-xylazine decreased time spent in wake and increased time in NREMS sleep in males. While no significant interaction effects were noted, both of these effects were concentrated in the dark period overall, mirroring the effect of fentanyl-only treatment. Notably, no changes in wake and NREMS were identified in females between groups despite a similar pattern of effect seen between fentanyl and fentanyl-xylazine as described before.

Opposingly, REMS was decreased in females treated with yohimbine compared to vehicle, producing an effect similar to fentanyl alone. And no effect was found in males despite a similar pattern seen in males given fentanyl and fentanyl-xylazine as described before.

### Yohimbine Reverses Fentanyl-Xylazine Wake and NREMS Bout Duration Alterations to Mirror Fentanyl in Females

To test if the effect of xylazine adulteration was dependent on the α_2_-adrenergic system we treated fentanyl-xylazine mice with yohimbine, an α_2_-adrenergic antagonist. Yohimbine has previously been shown to be therapeutic for multiple disorders including erectile dysfunction, metabolic and cardiovascular disease, and inflammation (PMID: 39684567). More recently, this α_2_-adrenergic antagonist exhibits efficacy in alleviating sleep apnea (PMID: 28239660). Therefore, we sought to determine whether yohimbine treatment would mitigate the sleep and respiratory effects of fentanyl-xylazine. Two-way repeated measures ANOVA revealed significant main effects of treatment (F (1, 6) = 11.43, P = 0.0148) and light-dark phase (F (1, 6) = 9.858, P = 0.0201) for average wake bout duration in females (Figure 6A). No significant differences were noted for the interaction between treatment x light-dark phase (F (1, 6) = 0.8546, P = 0.3909).

**Figure 6.**
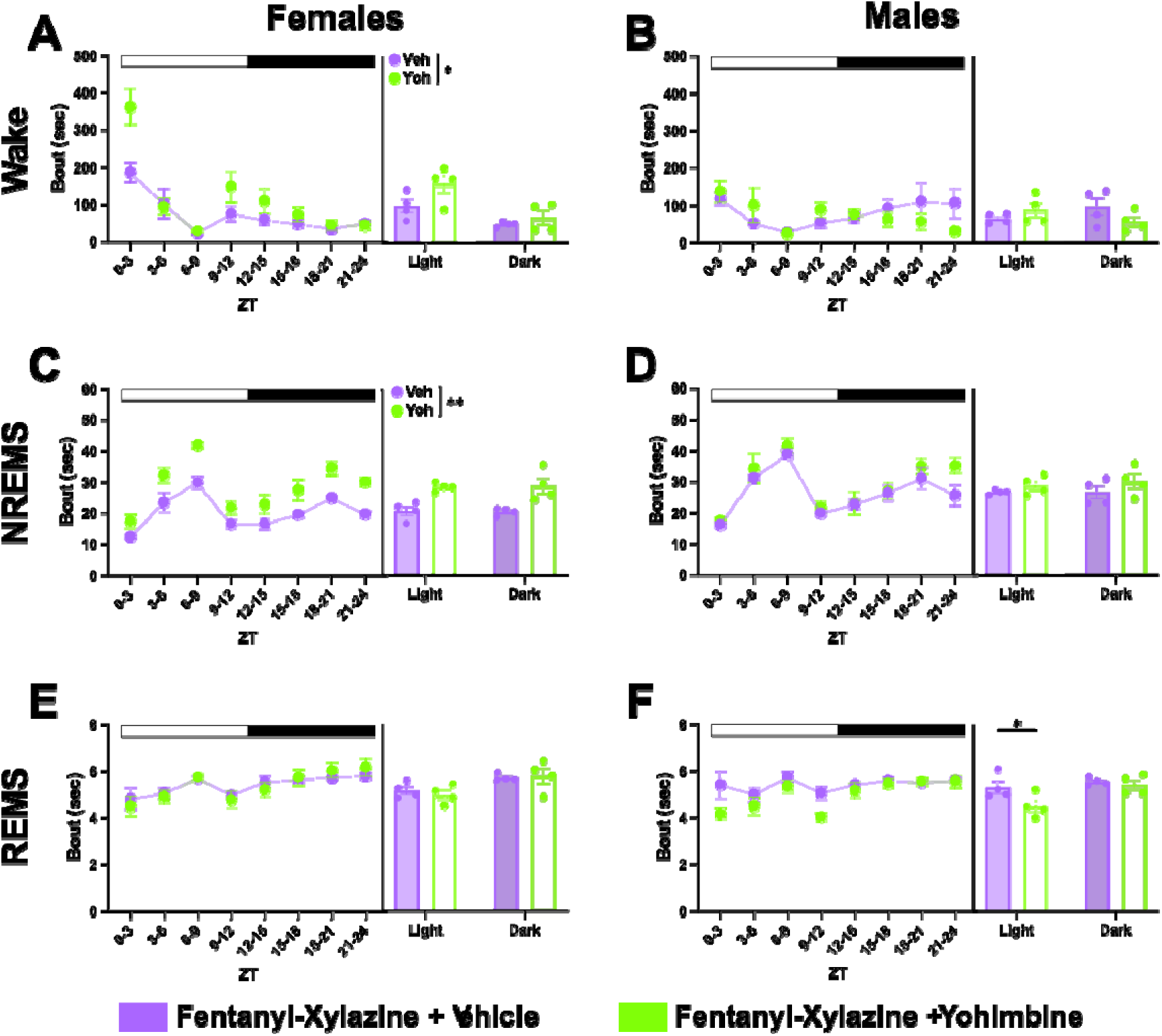
Alterations in bout durations across vigilance states from yohimbine treatment in female and male mice. **A** Average wake bout duration across seven days of recording expressed as in tri-ZT and light/dark bins in females. **B** Average wake bout duration across seven days of recording expressed as in tri-ZT and light/dark bins in males. **C** Average NREMS bout duration across seven days of recording expressed as in tri-ZT and light/dark bins in females. **D** Average NREMS bout duration across seven days of recording expressed as in tri-ZT and light/dark bins in males. **E** Average REMS bout duration across seven days of recording expressed as in tri-ZT and light/dark bins in females. **F** Average REMS bout duration across seven days of recording expressed as in tri-ZT and light/dark bins in males. Data presented as mean ± SEM; *p < 0.05, **p < 0.01.

Overall, yohimbine combined with fentanyl-xylazine treatment resulted in increased wake bout duration compared to vehicle in females. Looking per ZT indicates that this increase centered around injections akin to fentanyl-only treatment. For males, two-way repeated measures ANOVA indicated no significant main effects (treatment: F (1, 6) = 0.2018, P = 0.669, light-dark phase: F (1, 6) = 5.437e-005, P = 0.9944) or interaction effect (F (1, 6) = 2.904, P = 0.1392) (Figure 6B). Overall, yohimbine combined with fentanyl-xylazine had no effect on wake bout duration in males.

For average NREMS bout duration, a main effect of treatment was found in females (F (1, 6) = 26.86, P = 0.002), but not light-dark phase (F (1, 6) = 0.0003, P = 0.9867) or an interaction effect (F (1, 6) = 0.068, P = 0.803) (Figure 6C). Yohimbine combined with fentanyl-xylazine treatment decreased the average NREMS bout duration compared to vehicle across light-dark phase in females akin to fentanyl-only administration. No differences were found in males (treatment: F (1, 6) = 3.887, P = 0.0961, light-dark phase: F (1, 6) = 0.1101, P = 0.751, interaction: F (1, 6) = 0.1097, P = 0.7517) (Figure 6D).

In females, for average REMS bout duration, there was no significant main effect of treatment (F (1, 6) = 0.0151, P = 0.9061) nor interaction effect (F (1, 6) = 0.8292, P = 0.3976), only a significant main effect of light-dark phase (F (1, 6) = 20.93, P = 0.0038) (Figure 6E). As such, there were no changes in average REMS bout duration between yohimbine and vehicle-treated animals. In males, however, there was a significant main effect of light-dark phase (F (1, 6) = 17.60, P = 0.0057) and interaction effect (F (1, 6) = 6.331, P = 0.0455), but no significant main effect of treatment (F (1, 6) = 2.738, P = 0.1491) (Figure 6F). Holm corrected post-hoc testing indicated fentanyl-xylazine-yohimbine treated male mice had lower average REMS bout duration compared vehicle-treated mice during the light period (P = 0.0451). No differences were noted in the dark period.

In summary, yohimbine treatment increased average wake and NREMS bout duration—creating a profile more akin to fentanyl-only treatment—in female mice only. In male mice, yohimbine largely did not antagonize the effects of xylazine on sleep-wake bout duration and instead uniquely decreased average REMS bout duration during the light period.

### Baseline-Adjusted Behavioral Analysis Indicates No Significant Acute Spontaneous Withdrawal Phenotype in Fentanyl- or Fentanyl–Xylazine-Treated Mice

Females and males were video recorded prior to drug injections and approximately 18 hours after last injection and evaluated for signs of withdrawal (Figure 1). In females, linear mixed effects revealed a significant main effect of treatment ((F(3, 8) = 4.08, p = 0.0497) and interaction effect (F(3, 8) = 5.83, p = 0.0207) for shakes only (Figure 7). Holm post-hoc testing indicated that shakes were significantly decreased in xylazine and fentanyl treated mice compared to saline (P = 0.0409, P = 0.0368). No other behaviors were significantly different in females accounting for baseline. In males, linear mixed effects revealed a significant main effect of treatment (F(3, 16) = 3.64, p = 0.0355), but not an interaction effect (F(3, 16) = 0.96, p = 0.4349).

**Figure 7.**
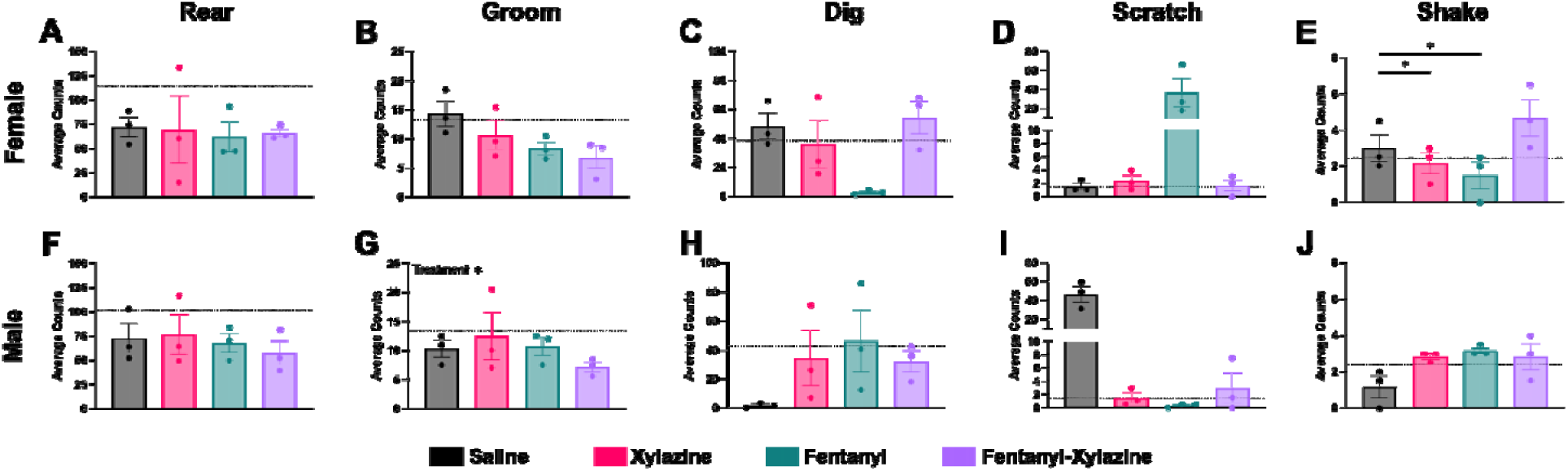
Acute spontaneous withdrawal behaviors following fentanyl and fentanyl-xylazine exposure in female and male mice. Behaviors manually scored from 15 min videos collected at baseline and during acute withdrawal from chronic fentanyl. **A-E** Average counts of withdrawal behaviors in females. **F-J** Average counts of withdrawal behaviors in males. Data presented as mean ± SEM; *p < 0.05.

### Xylazine Adulteration Enhances the Respiratory Depressive Effects of Fentanyl Primarily in Males

Naïve female and male mice were given escalating doses of fentanyl with or without a fixed dose of subanesthetic xylazine (Figure 1). Results represent only mice with a detectable depression in respiration (Females: 81.25%, Males: 84.38% of tested mice per sex, respectively). Supplemental Figure 1 shows a representative example of using piezoelectric sensors to isolate and capture distinct respiratory characteristics in mice.

In females, two-way ANOVA indicated a significant main effect of treatment (F (1, 18) = 5.699, P = 0.0282) and dose (F (3, 18) = 3.179, P = 0.0491), with no significant interaction (F (3, 18) = 1.656, P = 0.212) in respiratory depression (Figure 8A). This demonstrated that fentanyl-xylazine induced greater respiratory depression than fentanyl alone across doses, and that different doses were associated with varied levels of depression across treatments. Holm-corrected post-hoc comparisons of dose within the fentanyl-xylazine group indicated that 640 µg/kg fentanyl + xylazine produced greater depression than xylazine alone (P = 0.0485) and 1.28 mg/kg fentanyl + xylazine (P = 0.0178), but not 320 µg/kg + xylazine (P = 0.6149). No significant differences were detected among fentanyl-only doses.

**Figure 8.**
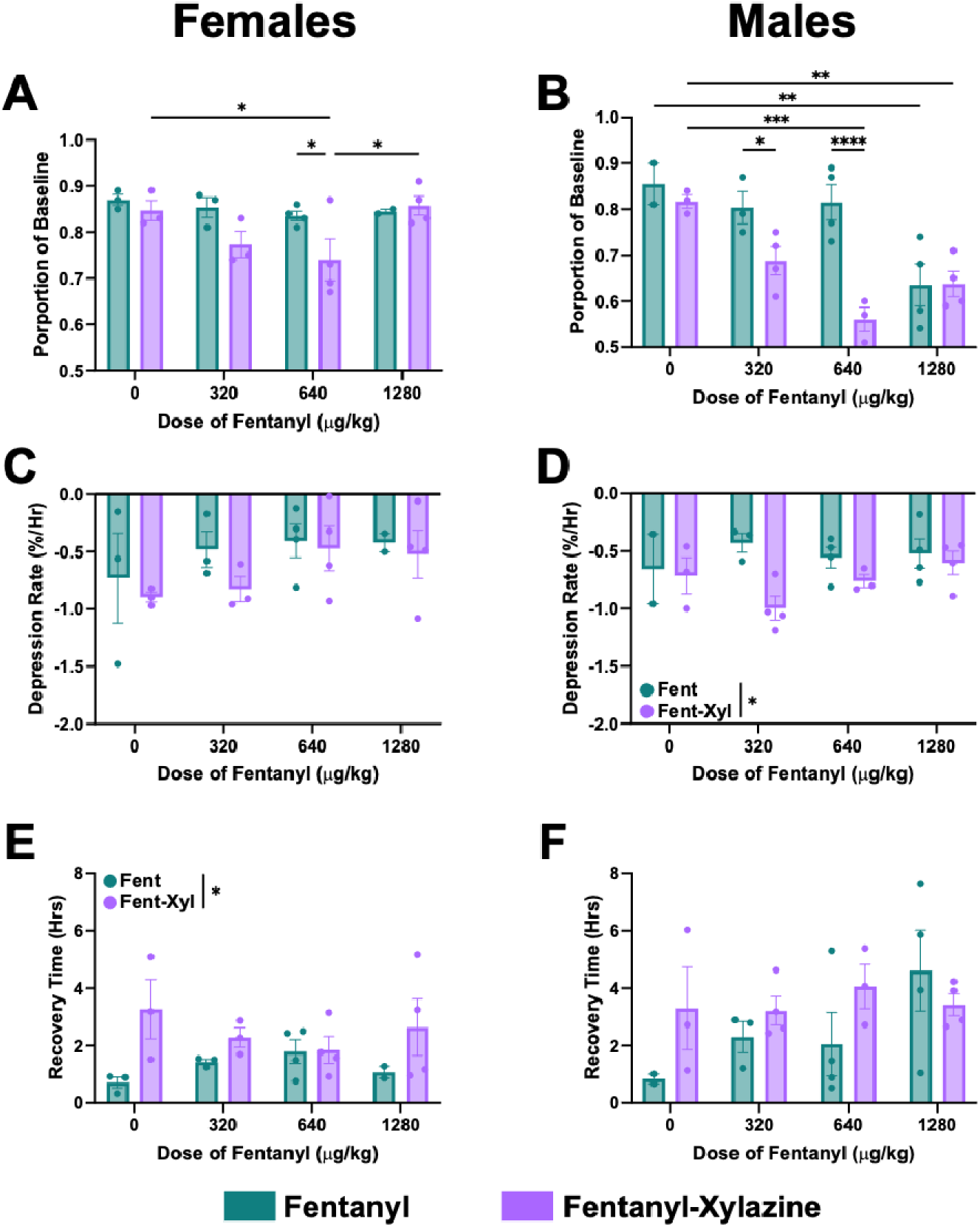
Evaluation of respiratory changes from escalating doses of fentanyl with and without a fixed dose of xylazine in female and male mice. **A** Proportion of baseline breathing in females. **B** Proportion of baseline breathing in males. **C** Rate of respiratory depression in females. **D** Rate of respiratory depression in males. **E** Time to recover back to baseline breathing rate in females. **F** Time to recover back to baseline breathing rate in males. Data presented as mean ± SEM; *p < 0.05, **p < 0.01, ***p < 0.001, ****p < 0.0001.

Conversely, in males, two-way ANOVA indicated significant main effects of treatment (F (1, 19) = 15.65, P = 0.0008), dose (F (3, 19) = 10.34, P = 0.0003), and a significant interaction (F (3, 19) = 5.295, P = 0.008) in respiratory depression (Figure 8B). Holm-corrected post-hoc comparisons showed that 320 µg/kg and 640 µg/kg fentanyl + xylazine doses were significantly lower than the same fentanyl only doses (P = 0.0308, P < 0.0001, respectively). At 1.28 mg/kg fentanyl, the addition of xylazine had no effect (P = 0.9572). No differences were detected between saline and xylazine controls. Notably, both 640 µg/kg and 1.28 mg/kg fentanyl + xylazine doses resulted in significantly more depressed respiration compared to xylazine-only treatment (P = 0.0007 and P = 0.0093, respectively). Among fentanyl-only doses, only 1.28 mg/kg fentanyl showed significantly more depressed respiration compared to saline (P = 0.0056).

Beyond total respiratory depression, there were also changes in respiratory depression rate and recovery time from respiratory depression. In females, there was no change in respiratory depression rate by treatment (F (1, 18) = 1.340, P = 0.2613), dose (F (3, 18) = 1.441, P = 0.2639), or interaction (F (3, 18) = 0.1829, P = 0.9066) (Figure 8C). However, for recovery time, a main effect of treatment only was noted (F (1, 18) = 7.36, P = 0.0143) (Figure 8E). The addition of xylazine ultimately increased recovery time in females. In males, the inverse was true, with a main effect of treatment only for respiratory depression rate (F (1, 19) = 6.761, P = 0.0176) (Figure 8D), but no change in recovery time by treatment (F (1, 19) = 2.188, P = 0.1555), dose (F (3, 19) = 1.314, P = 0.2989), or interaction (F (3, 19) = 1.425, P = 0.2666) (Figure 8F). The addition of xylazine decreased respiratory depression rate in males.

### Peripheral Molecular Analysis

#### Males and Females Do Not Develop Skin Lesions from Fentanyl-Xylazine Exposure and Show No Molecular Skin Integrity Abnormalities

No mice developed visible lesions or other skin abnormalities following fentanyl-xylazine treatment. Because gross examination may not detect more subtle or subclinical alterations in skin integrity, we collected skin biopsies at the end of the study to assess molecular markers associated with extracellular matrix remodeling and tissue structure. Specifically, we examined matrix metalloprotease 2 (Mmp2), which is involved in extracellular matrix degradation and remodeling, and fibronectin 1 (Fn1), an important extracellular matrix component involved in tissue structure and repair. Together, these markers allowed us to determine whether fentanyl-xylazine treatment was associated with molecular changes in the skin despite the absence of overt abnormalities. Two-way ANOVA revealed no effect of treatment (*Mmp2*: F (3, 41) = 1.426, P = 0.2489; *Fn1*: F (3, 41) = 0.3795, P = 0.7683), sex (*Mmp2*: F (1, 41) = 1.982, P = 0.1667; *Fn1*: F (1, 41) = 0.97, P = 0.33), or interaction (*Mmp2*: F (3, 41) = 2.341, P = 0.0874; *Fn1*: F (3, 41) = 0.1616, P = 0.9216) for both genes (Figure 9 A&B).

**Figure 9.**
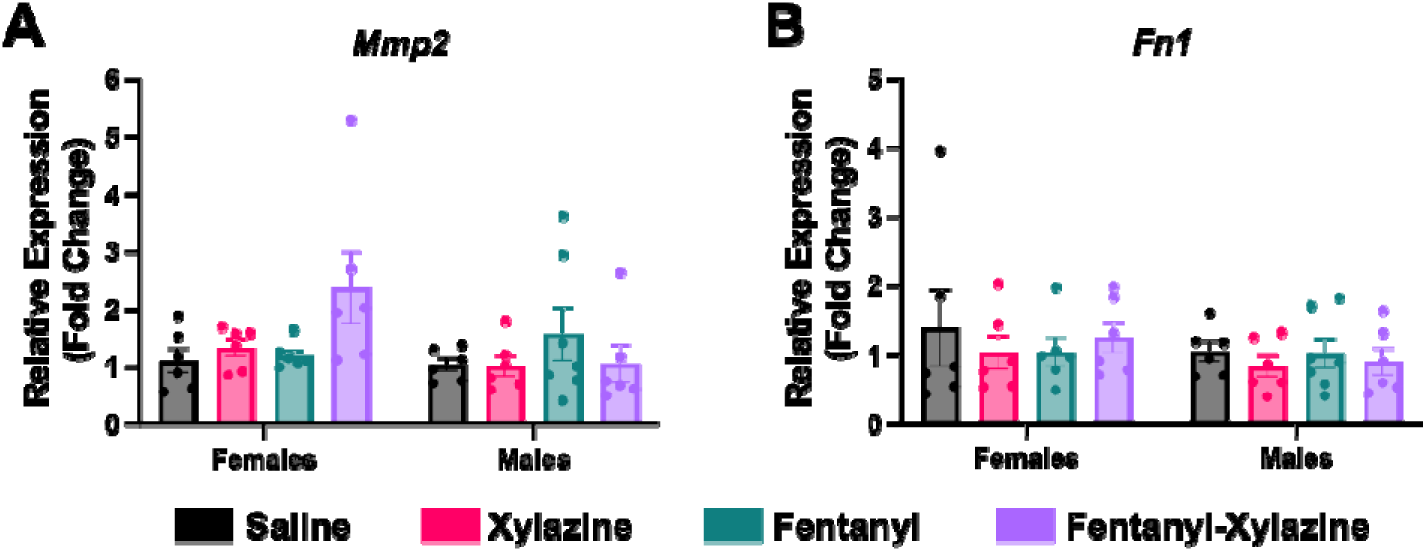
Molecular analysis of skin following fentanyl and fentanyl-xylazine exposure in males and females. **A** Expression of matrix metalloprotease protein 2 (*Mmp2*) by sex. **B** Expression of fibronectin 1 (*Fn1*) by sex. Data presented as mean ± SEM.

#### Fentanyl-Xylazine Elevated FGF21 in Mouse Serum

Serum was obtained from mice while drugs were on board and processed via targeted proteomics. Linear mixed-effects model was performed for sex in the Olink Target 48 assay (FDR corrected) and a main effect of treatment was identified for five proteins in females: FGF21 (F (3, 16.4) = 16.6, P < 0.0001), IL10 (F (3, 20) = 9.29, P = 0.00047), IL2 (F (3, 16.5) = 10.4, P = 0.00046), CCL4 (F (3, 16.4) = 8.09, P = 0.00156), and CXCL2 (F (3, 16.3) = 8.08, P = 0.00161) and two protein in males: CCL2 (F(3, 16.1 = 25.8, P< 0.0001), FGF21(F(3, 20.0) = 18.0, P< 0.0001). Tukey corrected post-hoc testing was then performed on all proteins to assess changes between drug groups. In females, FGF21 expression was significantly elevated due to xylazine (P = 0.0022) and fentanyl-xylazine (P < 0.0001) compared to saline. Moreover, fentanyl-xylazine had the higher expression compared to fentanyl (P = 0.0023) (Figure 10A). In males, FGF21 expression was significantly elevated due to xylazine (P = 0.00035) and fentanyl-xylazine (P < 0.0001) compared to saline and fentanyl-xylazine was also significantly higher than fentanyl-only (P = 0.0016) (Figure 10F).

**Figure 10.**
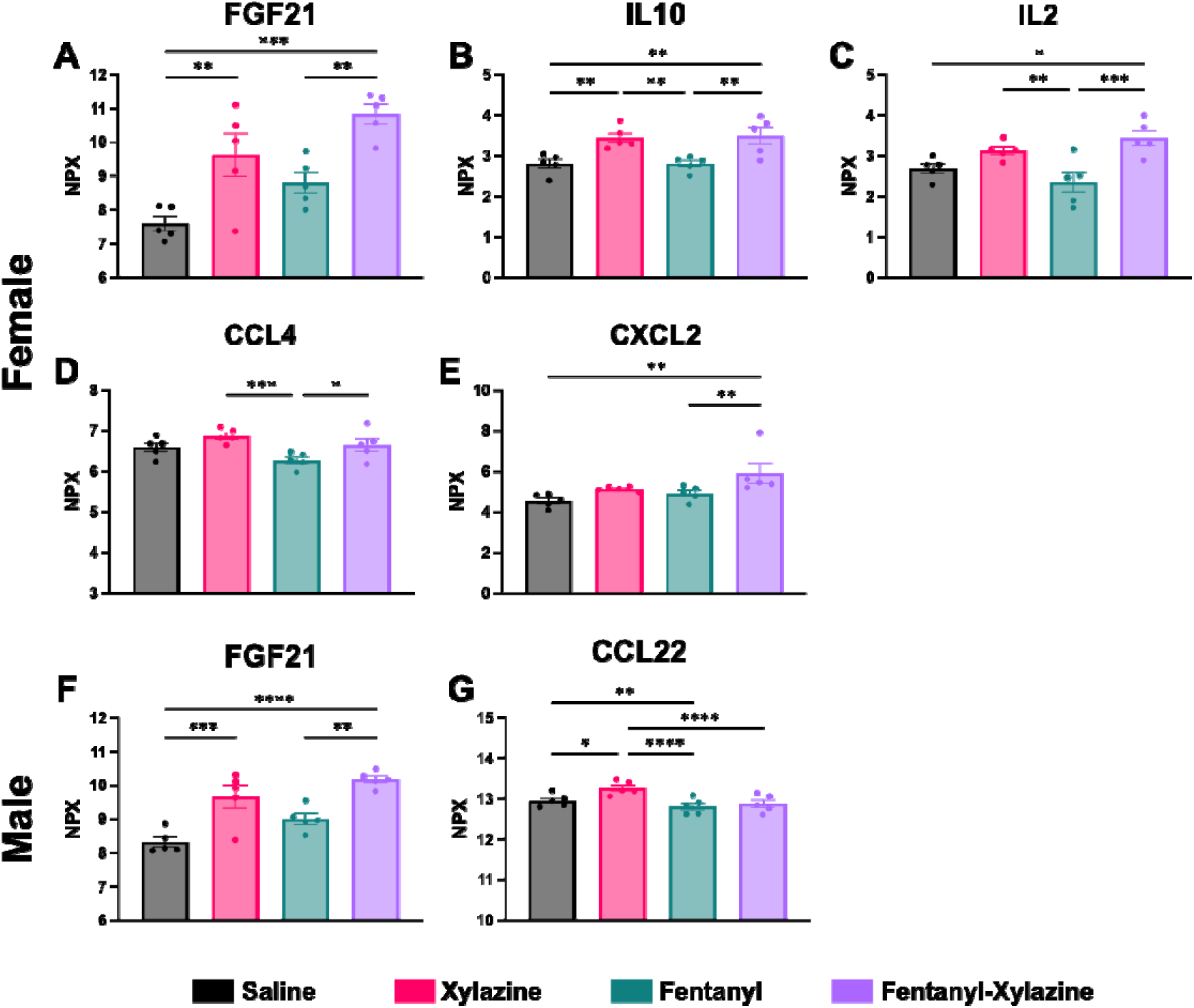
Targeted immune proteomic hits from serum of female and male mice on fentanyl and fentanyl-xylazine. **A** Normalized protein expression of fibroblast growth factor 21 (FGF21) per drug condition. **B** Normalized protein expression interleukin-10 (IL10) per drug condition. **C** Normalized protein expression of interleukin-2 (IL2) per drug condition. **D** Normalized protein expression of Chemokine (C-C motif) Ligand 4 (CCL4) per drug condition. **E** Normalized protein expression of C-X-C motif chemokine ligand 2 (CXCL2)22 (CCL22) per drug condition. **F** Normalized protein expression of fibroblast growth factor 21 (FGF21) per drug condition **G** Normalized protein expression of C-C motif chemokine ligand 22 (CCL22) per drug condition Data presented as mean ± SEM of NPX (normalized protein expression); *p < 0.05, **p < 0.01, ***p < 0.001, ****p < 0.0001.

For IL10 in females, xylazine and fentanyl-xylazine led to significantly elevated expression compared to saline (P = 0.0087, P = 0.0048) and fentanyl (P = 0.0094, P = 0.0051) but were not different from each other (Figure 10B). Similarly, IL2 was elevated the most in fentanyl-xylazine-treated females including saline (P = 0.0152) and fentanyl (P = 0.0004), but not different from xylazine. Xylazine trended in the same direction as fentanyl-xylazine but was only significantly elevated compared to fentanyl (P = 0.0010), and fentanyl was not different than saline (Figure 10C). CXCL2 was also significantly elevated in fentanyl-xylazine treated females compared to saline (P = 0.0014) and fentanyl (P = 0.0079), but not xylazine (Figure 10E). Lastly, CCL4 was significantly lower in fentanyl-treated females compared to xylazine and fentanyl-xylazine (P = 0.00086, P = 0.0265) but neither group was significantly different than saline (Figure 10D).

Furthermore, CCL2 in males, was significantly elevated in the xylazine group compared to saline (P = 0.00095), fentanyl (P < 0.0001), and fentanyl-xylazine (P < 0.0001). Additionally, fentanyl-only was significantly lower than saline (P = 0.0090), but not fentanyl-xylazine (Figure 10G).

## Discussion

Recent shifts in the fentanyl drug supply have led to significant adulteration with xylazine (D’Orazio et al., 2023). Emergency department reports have found that the combination of fentanyl and xylazine increases the risk for cardiac complications and coma due to opioid overdose (Love et al., 2023). Our current findings align with these clinical outcomes, as well as expand our understanding of how fentanyl-xylazine effects sleep-wake architecture, respiration, opioid withdrawal, and associated molecular changes. Specifically, we found that fentanyl administration in the light period increased wake time followed by an increase in NREMS in the dark as previously shown (Sharma et al., 2024). When xylazine was administered with fentanyl, this homeostatic response in the dark period was blunted in males despite equivalent changes in sleep disruption during the light period. This effect may partially be explained by changes in bout duration. Fentanyl-xylazine precluded the markedly decreased average wake bout duration associated with fentanyl-only which may have been protective against the need for homeostatic rebound sleep (Tietzel and Lack, 2001). However, it is worth noting that during this same period, fentanyl-xylazine mice also had the shortest average NREMS bout duration. In females, the fentanyl-xylazine administration largely compared to fentanyl; however, similar to males, xylazine with fentanyl normalized average wake bout duration and simultaneously reduced average NREMS bout duration. Overall, the addition of xylazine to fentanyl induced specific sleep-wake changes compared to fentanyl-only exposure.

Using yohimbine, we observed a sex-dependent alteration in the sleep–wake effects of fentanyl–xylazine administration that is consistent with a contribution of α - adrenergic receptor antagonism. In males, yohimbine treatment created an effect similar to fentanyl-only in percent time in wake and NREMS but not wake and NREMS bout duration. In females, however, yohimbine treatment produced an effect similar to fentanyl-only in wake and NREMS bout duration, but not percent time in wake and NREMS. The reason for this sex-specific difference is unclear, however, recent evidence indicates that xylazine displays kappa opioid activity and there are established, sex-specific differences in KOR expression in mice (Bedard et al., 2024; Chartoff and Mavrikaki, 2015).

Peripheral molecular analyses revealed elevated immune markers, including IL-10 and IL-2, in serum from fentanyl–xylazine-treated female mice compared with saline and fentanyl-only. This adds to previous studies highlighting that changes in immune markers including IL10 suggest a complex immune modulation pattern (Bryant et al., 2021; Mizher et al., 2020). IL10 is known as the primary anti-inflammatory cytokine, whereas IL2 is known for its role as T cell growth factor. Together they can work synergistically to enhance CD8+ T cell activity. Consistent with this possibility, Mazahery and colleagues reported altered CD8+ T cell phenotypes in individuals receiving chronic methadone treatment (Mazahery et al., 2020). Additionally, CXCL2 is a chemoattractant known to recruit opioid-containing neutrophils and stimulate opioid release through CXCR2 (Machelska, 2007). No changes were seen in splenic immune markers including IL10 (Supplemental Figure 2), suggesting that it is not the primary site of immune dysfunction due to fentanyl and fentanyl-xylazine exposure.

Confirming findings from previous studies, we showed that xylazine can enhance respiratory depression (Demery et al., 2025b). Interestingly, female mice were more resistant than male mice to fentanyl-induced respiratory depression, consistent with findings by Smith and colleagues, who observed a similar sex difference at the LD50 and LD80 doses of fentanyl (Smith et al., 2023). This suggests that there may be some sex-specific protective factors at least in mice. The highest dose of fentanyl and fentanyl-xylazine led to improved breathing in males and females which may reflect recruitment of hypoxia-evoked autoresuscitative gasping or other compensatory respiratory responses that emerge during profound hypoxia (Bush and Ramirez, 2024; Nuding et al., 2024; Orr et al., 2017).

Lastly, fentanyl-xylazine mice had significantly elevated levels of FGF21 predominantly in females, an important brain penetrant hepatokine for metabolic maintenance, that has been shown to reduce opioid preference in mice in at least one study (Dorval et al., 2022). Importantly, there is also a strong link between FGF21 and reduced alcohol consumption (Flippo et al., 2022; Liangpunsakul and Leggio, 2025; Wang et al., 2022). Genome-wide association studies have also found that FGF21’s receptor is associated with alcohol drinking, and several cross-species studies have found that higher levels of FGF21 curb alcohol drinking (Choi et al., 2023; Ho et al., 2022; Schumann et al., 2016; Wang et al., 2024). Other data have indicated that FGF21 may be a potential biomarker of poor sleep (Chen et al., 2025; Huang et al., 2023). Indeed, recent survey data suggest that individuals using fentanyl adulterated with xylazine report lower opioid consumption (Sibley et al., 2025) which is also true in rodent models of intravenous self-administration (Khatri et al., 2024; Sadek et al., 2023). Whether elevated FGF21 contributes to this pattern in humans remains unknown and warrants further investigation.

Several limitations should be considered when interpreting these findings. First, xylazine was administered at a single subanesthetic dose selected based on previous work (Khatri et al., 2024; Sadek et al., 2023) to model adulterant-level exposure. Because α_2_-adrenergic and putative kappa-opioid effects of xylazine are dose-dependent, a broader xylazine dose-range and additional fentanyl doses in the chronic sleep and withdrawal arms would help establish the dose-dependence of the interactions reported here. Second, there are no yohimbine-only control mice and while yohimbine is primarily an α_2_-AR antagonist it also has activity at serotonin and dopamine receptors. Third, our molecular characterization was restricted to peripheral serum and spleen together with skin and did not include central nervous system tissue; the sleep-wake and respiratory phenotypes therefore cannot yet be mapped onto specific neural substrates, and the sex-dependent yohimbine effects remain associative rather than mechanistically resolved. Fourth, the elevation of FGF21 is correlational; whether it contributes to the reduced opioid consumption reported in people using xylazine-adulterated fentanyl, or instead reflects hepatic or metabolic stress, cannot be distinguished from these data. Finally, respiratory measurements were derived from piezoelectric recordings rather than plethysmography, and while this approach captured treatment- and sex-dependent differences in respiratory depression, it does not resolve tidal volume or blood-gas endpoints. These constraints define clear priorities for future mechanistic and translational work.

## Conclusions

This study provides a detailed preclinical characterization of the combined effects of fentanyl and xylazine, revealing unique, sex-specific alterations in sleep-wake dynamics and enhanced risk of respiratory depression. Notably, despite widespread observations of necrotic lesions in humans in response to combined fentanyl-xylazine use, we did not observe similar skin lesions in our mouse model. Molecular analyses demonstrated peripheral metabolic changes associated with fentanyl-xylazine exposure known to reduce alcohol use and sleep disruption, which warrant future studies to ascertain if they are involved in reducing opioid use and withdrawal. Overall, these results underscore the value of polysubstance investigations, which remain uncommon even in the preclinical literature, and highlight the need to clarify the mechanisms underlying opioid co-use, its impact on overdose risk, and its potential sex-dependent effects.

## CRediT Authorship Contribution Statement

**Mackenzie C. Gamble:** Conceptualization, Methodology, Data Curation, Formal analysis, Investigation, Data Curation, Visualization, Writing - Original Draft, Writing - Review & Editing, Visualization, Project Administration**, Samara J. Vilca:** Methodology, Data Curation, Formal Analysis, Project Administration, Writing - Review & Editing**, Benjamin R. Williams:** Methodology, Data Curation, Formal analysis, **Grace Guidon:** Data Curation**, Vivian Dang:** Data Curation**, Ryan W. Logan:** Conceptualization, Methodology, Project Administration, Supervision, Writing - Review & Editing, Funding Acquisition, Resources.

## Funding

Work was supported by National Institutes of Health: NIDA R01DA061243 (RWL and ZF).

## Declaration of Competing Interests

The authors declare no potential conflicts of interest.

## Notes

### Competing Interest Statement

The authors have declared no competing interest.

